# Stress fibers are embedded in a contractile cortical network

**DOI:** 10.1101/2020.02.11.944579

**Authors:** Timothée Vignaud, Calina Copos, Christophe Leterrier, Qingzong Tseng, Laurent Blanchoin, Alex Mogilner, Manuel Théry, Laetitia Kurzawa

## Abstract

Contractile actomyosin networks generate intracellular forces essential for the regulation of cell shape, migration, and cell-fate decisions, ultimately leading to the remodeling and patterning of tissues. Although actin filaments aligned in bundles represent the main source of traction-force production in adherent cells, there is increasing evidence that these bundles form interconnected and interconvertible structures with the rest of the intracellular actin network. In this study, we explored how these bundles are connected to the surrounding cortical network and the mechanical impact of these interconnected structures on the production and distribution of traction forces on the extracellular matrix and throughout the cell. By using a combination of hydrogel micropatterning, traction-force microscopy and laser photoablation, we measured the relaxation of the cellular traction field in response to local photoablations at various positions within the cell. Our experimental results and modeling of the mechanical response of the network revealed that bundles were fully embedded along their entire length in a continuous and contractile network of cortical filaments. Moreover, the propagation of the contraction of these bundles throughout the entire cell was dependent on this embedding. In addition, these bundles appeared to originate from the alignment and coalescence of thin and unattached cortical actin filaments from the surrounding mesh.

## INTRODUCTION

Contractile forces play a key role in the regulation of cell and tissue morphogenesis (Heisenberg and Bellaïche, 2013; Murrell et al., 2015). At the level of tissues, they drive the shape changes supporting tissue remodeling such as extension (Rauzi et al., 2008), compaction (Maître et al., 2015) and folding (Hughes et al., 2018). At the level of the cell, they affect shape (Mogilner and Keren, 2009), migration (Leal-Egaña et al., 2017; Maiuri et al., 2015), division (Sedzinski et al., 2011) and differentiation (Kilian et al., 2010; McBeath et al., 2004). Contractile forces are produced mainly by actomyosin bundles or stress fibers in adherent cells (Chrzanowska-Wodnicka and Burridge, 1996; Katoh et al., 1998; Naumanen et al., 2008), and by a cortical meshwork of randomly oriented filaments in poorly-adherent cells (Chugh and Paluch, 2018; Chugh et al., 2017). To produce effective morphological changes, the magnitude and spatial distribution of contractile forces must be regulated and integrated over distances that are much longer than individual filaments (Agarwal and Zaidel-Bar, 2019; Murrell et al., 2015). However, the mechanism regulating the production and transmission of local forces throughout the cell is still poorly understood (Burridge and Guilluy, 2016; Kurzawa et al., 2017; Livne and Geiger, 2016). The progress in understanding this integration process has notably been limited by the technical challenges to manipulate the network locally while simultaneously measuring the impact on force production at the level of the entire cell.

Stress fibers are formed by the interaction and merging of pre-existing radial fibers and transverse arcs (Hotulainen and Lappalainen, 2006; Schulze et al., 2014; Tojkander et al., 2012, 2015). Transverse arcs are formed by the alignement and compaction of filaments at the cell front, as they are pulled by the actin network retrograde flow against cell anchorages (Burnette et al., 2011; Shemesh et al., 2009). As a result, actomyosin networks are composed of interconnected contractile elements that span the entire cytoplasm and serve as a template to transmit mechanical forces over long cellular distances (Cai et al., 2010; Hu et al., 2003; Wang et al., 1993). Laser photoablation experiments have indeed demonstrated that the photoablation of a single stress fiber could compromise the entire traction force field (Kumar et al., 2006; Tanner et al., 2010) and lead to variations in tension in all focal adhesions including those that are not at the ends of the ablated fibers (Chang and Kumar, 2013). Similarly, stretching cells unidirectionally can lead to tension increase in all focal adhesions whatever their orientation (Kumar et al., 2019). Hence, directional forces along specific actomyosin bundles can propagate to other bundles with which they are inter-connected. As a consequence, the tension in a stress fiber does not only depend on forces produced in that fiber but also on the connection and orientation of adjacent fibers (Kassianidou et al., 2017). This high degree of connection between actomyosin bundles can provide the mechanical coherence at the level of the cell (Chapin et al., 2012; Rossier et al., 2010; Smith et al., 2010). However, it is yet unclear how forces are transmitted from one stress fiber to the other.

Theoretical models of contractile networks have proposed two main paradigms to capture the mechanisms of force production and transmission in cells. In one paradigm, discrete models, that include high level details on the structure of the network, offer an accurate description of the stress fiber as a load-bearing structure and of the traction forces exerted on its anchorages to the extra-cellular matrix (Besser et al., 2011; Guthardt Torres et al., 2012; Kassianidou and Kumar, 2015; Luo et al., 2008; Stachowiak and Shaughnessy, 2009). These models are successful at providing a description in fine details of local force production, but fail to provide a global description of the traction-force field. In the other paradigm, continuous models provide a more global view of the contractile networks by incorporating only a few coarse-grained biophysical parameters and no particular structures such as stress fibers. Such models include those based on active gel theory, which describe the actomyosin network as a homogeneous compressible viscoelastic gel under boundary conditions. These models work well for describing force variations with changes in cell size and shape (Bischofs et al., 2008; Linsmeier et al., 2016; Oakes et al., 2014). Interestingly, elastic theory is even more accurate when the actual organization of actomyosin bundles and the spatial distribution of focal adhesions are taken into account (Von Erlach et al., 2018). These considerations suggest that the limitations of the discrete and the continuous models could be overcome by developing an intermediate model which takes into account both features of the network. It would help at fully capturing the global integration of mechanical forces in cells.

In this study, we used a unique combination of micropatterning, local photoablation of contractile elements, global-force measurements, and theoretical modeling to describe finely the mechanics of the subcellular actin networks. Our results revealed that contractile bundles were not only interconnected but also embedded in a continuous meshwork of cortical filaments along their entire length. This cortical meshwork not only transmitted bundle contraction throughout the cell but also contributed a significant part to the total force a cell exerted on its micro-environment.

## RESULTS

### Cells with prominent stress fibers produce a large amount of contractile force

We first tested experimentally if the presence of actin bundles could impact the magnitude and distribution of traction forces as compared to a more homogeneous network of the same size and shape (Linsmeier et al., 2016; Oakes et al., 2014). To that end, cells were plated on either dumbbell-shaped or pill-shaped micropatterns fabricated on hydrogel. The dumbbell-shaped micropattern was designed to promote in the cell the assembly of two stress fibers spanning the non-adhesive zone as previously described (Mandal et al., 2014; Théry et al., 2006), whereas the pill-shaped micropattern was designed to promote a more homogeneous network of actin filaments (Figure 1a). Both micropatterns were devised to promote the cells adopting the same shape.

**Figure 1.**
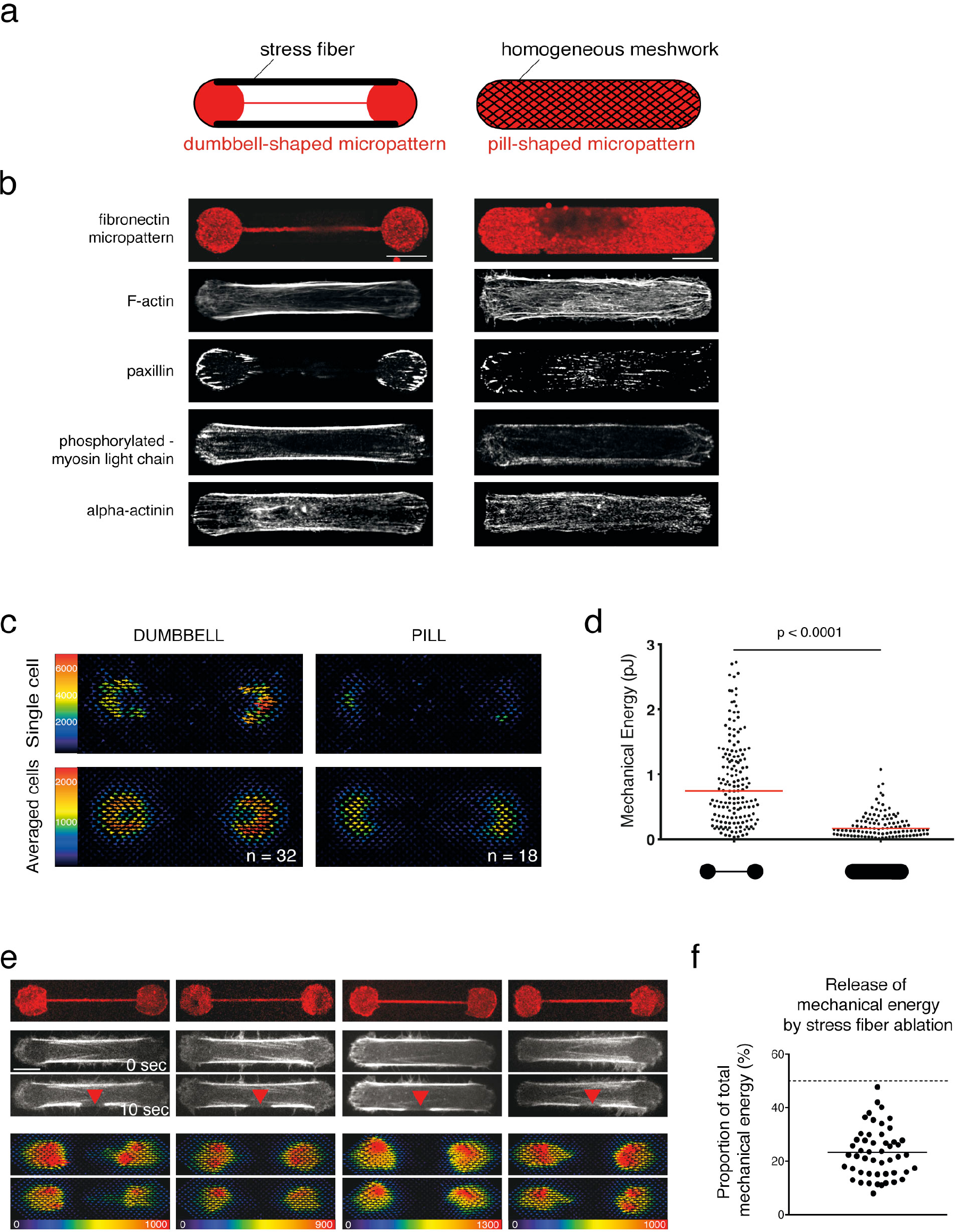
The stress fiber sets the magnitude of the traction force exerted by the cell but remains under tension after photoablation. a. Micropattern designs (59 μm length) and their respective outcomes in actin-network architecture. Dumbbell shape (left): Actin stress fibers (thick black lines) form between the two adhesive disks (red). Pill shape (right): Formation of a continuous actin mesh. b. Immunostainings of RPE1 cells spread on dumbbell-shaped (left panel) and pill-shaped polyacrylamide micropatterns (right panel), respectively. For each shape, single examples of representative cells are displayed. From top to bottom: micropatterns labeling (fibrinogen-Cy5); actin (phalloidin ATO-488); paxillin (Alexa-488); phosphorylated-Myosin light chain (CY3); alpha-actinin (CY3). Image scale bar = 10 μm. c. Traction-force maps of cells spread on dumbbell (left column) and pill micropatterns (right column) of 37 μm, respectively. Upper images display traction-force maps of single representative cells (scale bar in Pa). Lower images show averaged traction-force maps of cells. d. Scatter plot of the mechanical energies of single cells and associated p-value (Mann-Whitney t-test) (Dumbbell shape n=160, pill shape n =107). e. Force relaxation study upon peripheral stress-fiber photoablation. Left panel from top to bottom: Micropattern labeling (Fibrinogen-CY5); actin (LifeAct-GFP) before photoablation (0 sec) and after photoablation (10 sec, red arrow); corresponding traction-force maps of the initial forces; and traction-force maps after photoablation. Image scale bar = 10 μm. Force scale bar in Pa. f. Scatter plot of individual released mechanical energies after stress-fiber photoablation (% of the initial mechanical energy) for n = 49 cells.

As expected, two main peripheral stress fibers and only a few smaller and thinner internal bundles were formed in cells plated on dumbbell-shaped micropatterns (see representative example in Figure 1b, left panel, and an averaged image of several cells in Supplementary Figure 1, left panel). These structures concentrated crosslinkers of actin filaments and myosins, which are characteristic of stress fibers (Peterson et al., 2004). By contrast, numerous, more evenly distributed but thinner bundles of actin filaments were formed in cells on pill-shaped micropatterns (Figure 1b, right panels and Supplementary Figure 1, right panel).

The contractility generated by these cells forming two distinct cytoskeletal networks enclosed in a similar envelope was measured using traction force microscopy (TFM). As illustrated by the averaged traction-force maps and quantified by the mechanical energy that was transferred to the hydrogel, significantly higher mechanical energy was generated by cells containing the stress fibers (dumbbell micropattern) than the cells without (pill micropattern; Figure 1c, d). This result indicated that the organization of actomyosin components into stress fibers plays a major role in setting the magnitude of force a cell could generate and transmit to the substrate. However, these global force measurements could not reveal the quantitative contribution of individual stress fibers to the total force produced by the cells.

### Stress fibers are connected to surrounding actin cytoskeletal elements

By combining photoablation of the peripheral stress fibers with traction-force measurements, we assessed the specific contribution of these fibers to the global mechanical energy produced by cells plated on dumbbell micropatterns. To visualize the stress fibers, cells expressing either actin-GFP or LifeAct-GFP were used. A stress fiber was severed mid-length by localized pulsed-laser photoablation at 355 nm (see Supplementary Movie S1), and the release in fiber tension was captured by the relaxation in the hydrogel substrate (Figure 1e and Supplementary Movie S2). Surprisingly, the released energy from the cut of one of the two peripheral stress fibers was about 25 % of the total mechanical energy generated by the cell (Figure 1e, f), and was substantially lower than the expected 50% assuming the two peripheral stress fibers generate most of the mechanical energy.

This prompted us to investigate in more detail the relaxation of the severed stress fiber. Marks were photobleached along the fiber to monitor the entire relaxation pattern after the severing, (Colombelli et al., 2009) (Figure 2a). As previously described, the retraction of a severed end was characteristic of a visco-elastic relaxation (Koonce et al., 1982; Kumar et al., 2006; Strahs and Berns, 1979) (Figure 2b). Indeed, the parts of the fiber distal to the photoablation displayed minimal if any relaxation (Figure 2b, 2c), and contrasted with what would have been be expected with a stress fiber in isolation, in that the relaxation should be independent of the distance from the photoablation (Figure 2c). Hence, this result suggested that the fiber was not isolated but connected along its length to force-bearing elements which resisted the deformation when the fiber was severed. Similar observations have been made elsewhere in cells spread on uniform extra-cellular matrix coating (Colombelli et al., 2009). In this work, stress fibers appeared to be connected to the extra-cellular matrix via focal adhesion-like complexes which acted as elastic linkers, and were shown to accumulate zyxin at their severed ends following photoablation at locations where new adhesions were formed. However, with the dumbbell micropattern, stress fibers were above a non-adhesive substrate, precluding the possibility of transmembrane adhesions forming at the severed ends. This led us to hypothesize that peripheral stress fibers were not connected to the extra-cellular environment but to cortical actin filaments.

**Figure 2.**
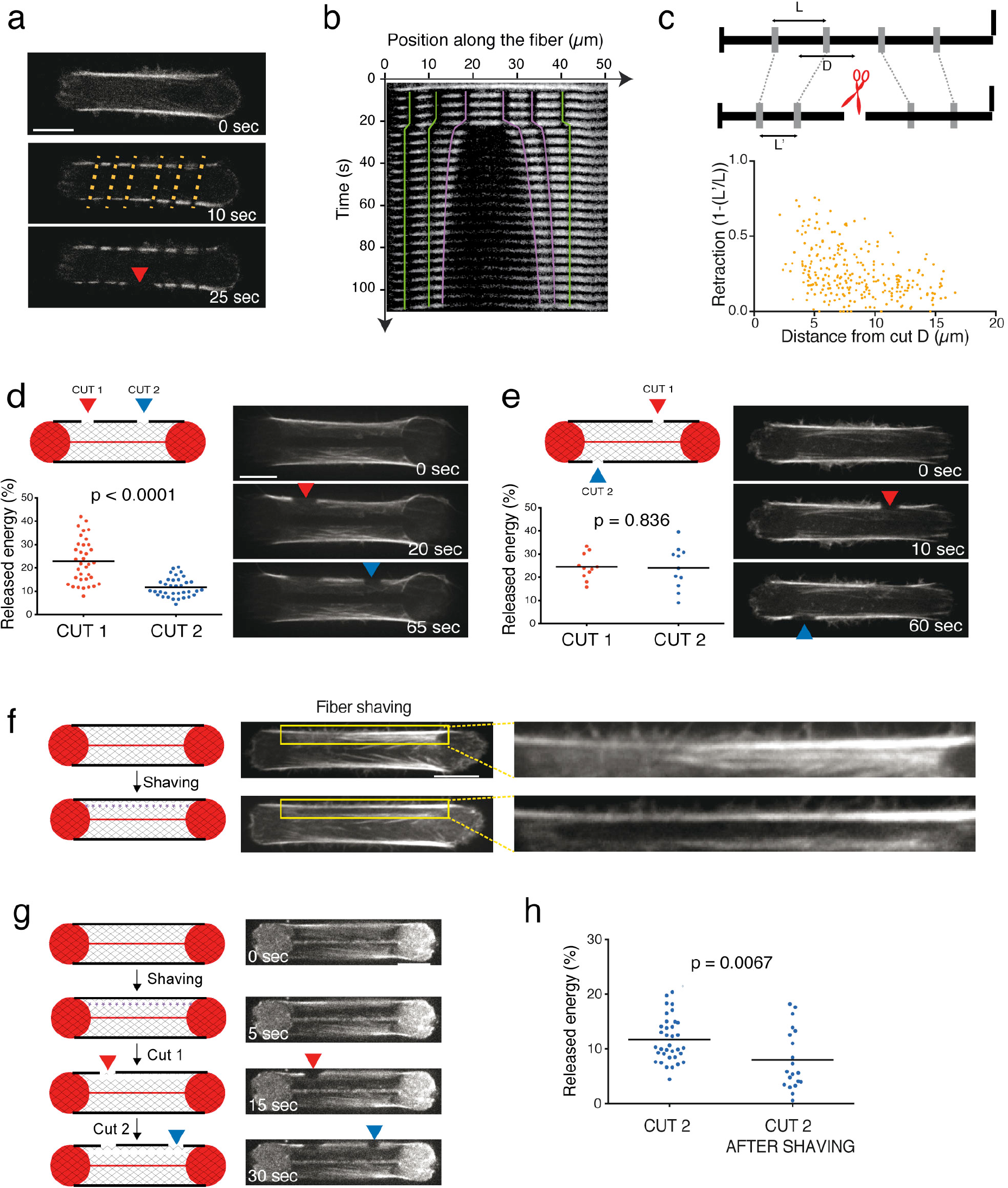
Stress fibers are connected to the surrounding actin cytoskeletal network and the connections impact the distribution of forces along the stress-fiber length and throughout the cell. a. From top to bottom: RPE1-actin-GFP time-lapse images. Stripes were added at regular intervals along the stress fiber by targeted photobleaching (at 10 sec, orange dashed lines); photoablation was then performed at the center of the stress fiber (at 25 sec, red arrow). Image scale bar = 10 μm. b. Corresponding kymograph with colored lines highlighting the retraction of the photobleached marks. c. Normalized retraction distance was calculated by dividing the length of each retracted segment (between two marks) by its initial length. Corresponding retraction values were plotted as a function of the initial distance of the photobleached segment from the photoablation site for n = 376 segments (58 cells analyzed). d. Scheme and corresponding images illustrating the experimental procedure of two sequential photoablations (cuts) on the same stress fiber at distinct locations (Cut 1=red arrow; Cut 2=blue arrow). Right panel displays time-lapse images of RPE1-LifeAct-GFP cells after the two sequential photoablations. Scatter plots of the released mechanical energy (percentage of the mechanical energy before photoablation) for the two types of photoablation sequence (n=35 cells). The p-value from a paired t-test is indicated on the plot. Image scale bar = 10 μm. e. Scheme and corresponding images illustrating the experimental procedure of two sequential photoablations (cuts) on the two stress fibers in the cell (Cut 1=red arrow; Cut 2=blue arrow). Right panel displays time-lapse images of RPE1-LifeAct-GFP cells after 2 photoablations. Scatter plots of the released mechanical energy (percentage of the mechanical energy before photoablation) for the two types of photoablation protocols (n=11 cells). The p-value from a paired t-test is indicated on the plot. f. Scheme and corresponding images of RPE1 expressing LifeAct-GFP illustrating stress-fiber shaving procedure along the stress fiber (purple dashed line), i.e. photoablation of a narrow region medial and parallel to the length of the fiber above the non-adhesive substrate on the hydrogel. The shaved region (dark area) sitting next to the stress fiber was highlighted in the yellow inset and corresponding zoomed images. Image scale bar = 10 μm. g. Left panels: Scheme illustrating the procedure of shaving (purple dashed line) followed by two sequential photoablations (cuts) on the adjacent peripheral stress fiber (Cut 1=red arrow; Cut 2=blue arrow). Middle panel: RPE1 cells labeled with SiR-actin and their corresponding micropatterns (fibrinogen-CY5) in a time-sequence corresponding to the shaving (T=5 sec), the first photoablation (T=15 sec, red arrow) and the second photoablation (T=30 sec, blue arrow). Image scale bar = 10 μm. h. Scatter plot of the mechanical energy released by the second photoablation of the stress fiber (percentage of the initial mechanical energy), alone (Cut 2; n=35 cells) or preceded by a shaving (Cut 2 after shav; n=28 cells). The p-value of Mann-Whitney t-test is indicated on the plot.

### Stress fibers are still under tension following laser photoablation

Importantly, and in contrast to the classical view of stress fibers pulling on focal adhesions only, the stress-fiber connection to adjacent actin cytoskeletal elements implied that photoablation should redistribute the tension of the fiber to the surrounding meshwork. As a result, the remaining parts of the severed fiber should still be under tension. To test this prediction, two photoablations were performed sequentially less than a minute apart on the same fiber. In confirmation of the prediction, a noticeable release of energy was associated with the second photoablation (Figure 2d). However, the amount of energy that was released by the second photoablation was much lower than the first one because the severed fiber had already relaxed and lost some of its elastic energy. To investigate whether this lower amount of energy release resulted from a non-specific injury to the cell due to the photoablations, the second photoablation was performed on the other intact fiber. In that case, approximately the same amount of energy was released as that after the first photoablation showing that the first photoablation did not impact cell integrity (Figure 2e).

Previous work has suggested that discrete connections of a peripheral stress fiber to other internal fibers can affect its relaxation pattern after severing (Kassianidou et al., 2017, 2019). However, the incomplete relaxation pattern of the severed peripheral stress fibers we observed here was not systematically associated with interconnections with internal fibers, as illustrated by the absence of visible fibers connected to the peripheral stress fibers in Figure 2a. This led us to hypothesize that a stress fiber was connected to a low-density and
widespread actin meshwork. This hypothesis was addressed by disconnecting the stress fiber from the meshwork through photoablation of a narrow region medial and parallel to the length of the fiber above the non-adhesive substrate on the hydrogel (a process we termed fiber shaving; Figure 2f and Supplementary Movie S3). With two successive photoablations of the stress fiber, the release of energy after the second photoablation was significantly lower when the fiber had been shaved first (Figure 2g, h), supporting the hypothesis that low-density fibers connections to the peripheral stress fiber prevented its complete relaxation after it was severed.

### Modeling the stress fibers embedding in an elastic meshwork

To investigate the properties of such a network of actin fibers embedded in a cortical meshwork, we built on the ideas developed in (Besser et al., 2011; Colombelli et al., 2009) to create a new biophysical model. In this model, and in contrast with previous ones, the stress fibers were not connected to the extra-cellular matrix but to the adjacent cortical meshwork. The cortical meshwork was described as a two-dimensional (2D) ensemble of elastic links connected by nodes. The stress fibers were modeled as elastic cables under tension and connected in series. The stress fibers were connected uniformly along their length to the cortical meshwork by elastic links (Figure 3a; described in detail in Supplementary Methods). The spring constants of the fiber-mesh and intramesh links were the same. The constraints on the model parameters are discussed in more detail in the Supplementary Methods.

**Figure 3.**
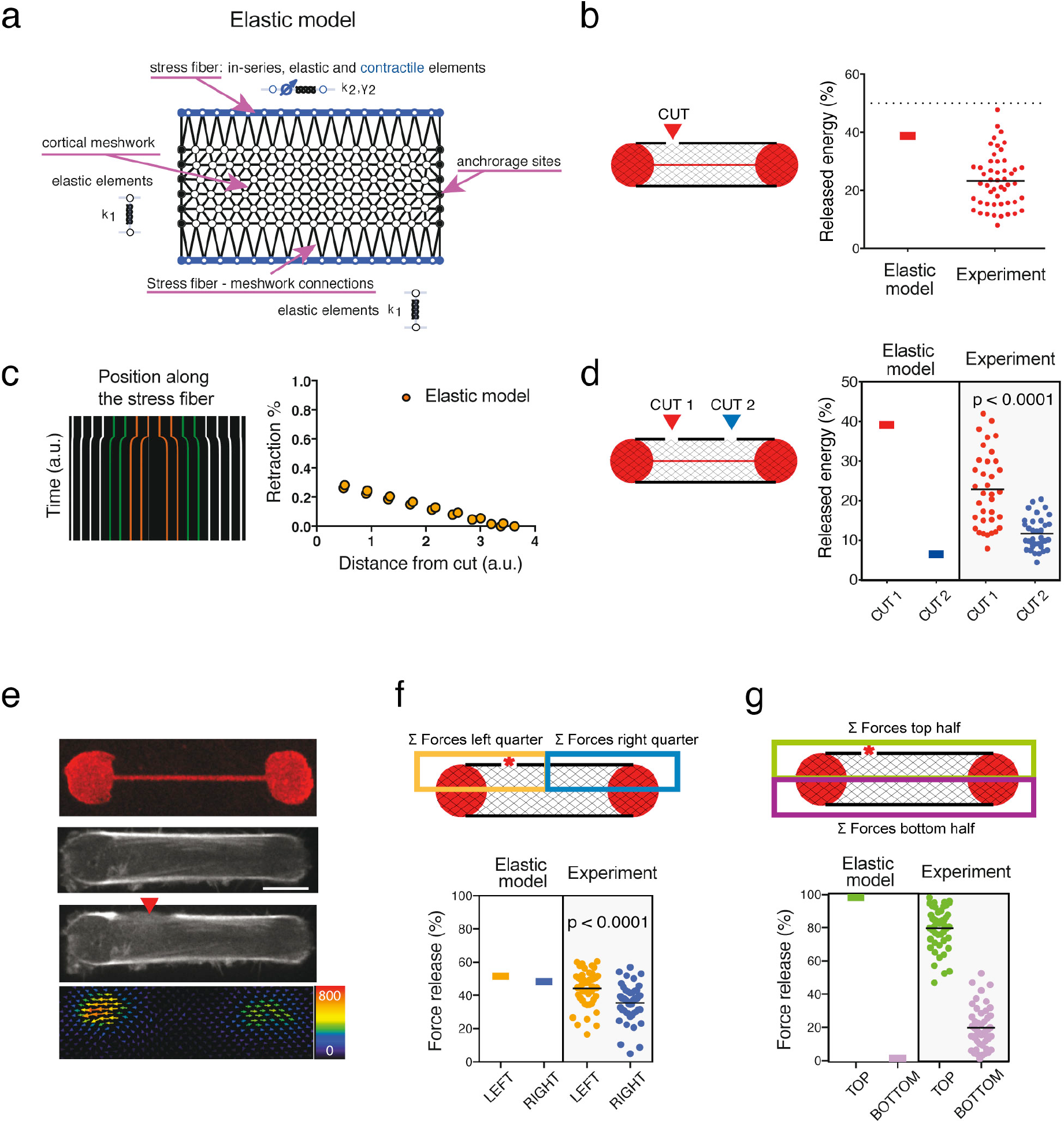
A model with active contractile stress fibers embedded in an elastic central mesh recapitulates key experimental findings of stress-fiber photoablation. a. Diagram illustrating all components in the model, including the isotropic elastic cortical mesh, contractile elastic fibers along the long edges of the mesh, and adhesive sites located along the short edges of the mesh. b. A single photoablation was reproduced in the model by a local reduction of the elastic stiffness in one link of the fiber. Plot displaying simulated traction-force loss as a result of the stress fiber ablation and corresponding experimental data. c. Simulated kymograph of a stress fiber indicating the movement of the regularly-spaced markers after simulated photoablation. Plot of the associated retraction percentage of these markers as a function of the distance from the ablation site. d. Scheme and plot displaying the predicted and experimental mechanical energy release after two sequential photoablations on the same stress fiber. The p-value from a paired t-test is indicated on the plot. e. Representative image of the dumbbell-shaped micropattern (fibrinogen-CY5) and RPE1-LifeAct-GFP cells displaying photoablation at a lateral side of the stress fiber (red arrow) and the associated relaxation traction force field after the photoablation. Image scale bar = 10 μm. Force scale bar in Pa. f. Spatial distribution of force relaxation along the stress fiber after stress-fiber photoablation (red star). Left panel: The release of traction forces was considered in partitioned zones of the cell, where the orange zone included half the stress fiber and the off-centered photoablation site, and the blue zone included the other half of the stress fiber. Right panel: Plot displaying the prediction of the model and the experimental measurements (n=47 cells) for the magnitude of released forces (as a percentage of the total force release) with respect to these zones. The p-value from the paired t-test is indicated on the plot. g. Spatial distribution of force relaxation across the cell after stress-fiber photoablation (red star). Left panel: The release of traction forces was considered in partitioned zones of the cell, where the green zone included the stress fiber with photoablation site, and the purple zone included the stress fiber without photoablation. Right panel: Plot displaying the prediction of the model and the experimental measurements (n=47 cells) for the magnitude of released forces (as a percentage of the total force release) with respect to these zones.

We first tested the response of our model to fiber severing by locally reducing the stiffness by 90% in one of the stress-fiber elements (Figure 3b). As in the experimental set up, mechanical energy was released but was significantly less than 50%, even though in the model, it was exclusively produced by the two peripheral fibers (Figure 3b). The model also accurately accounted for the limited retraction of the fiber at distal points from the site of the simulated photoablation (Figure 3c). Furthermore, the model captured the additional release of energy after a second photoablation of the same fiber (Figure 3d).

Other experimental observations prompted further investigations of the model. In the experimental set up, the release in tension was not equivalent on the two severed parts of the stress fiber, and was not restricted to its distal ends: rather, the release in tension appeared higher at the end that was closer to the photoablation and extended to the other side of the cell (Figure 3e and other examples in Supplementary Figure 2). By contrast, an isolated stress fiber would be expected to release the same amount of tension at both ends. This suggested that the cortical meshwork had an impact not only on the magnitude but also on the spatial distribution of traction-force production. To quantify this spatial effect in the model, the cell area was partitioned into four quadrants and the relative traction-force loss was measured in each quadrant as a percent amount of total traction-force loss in the cell (Figure 3f, g). Intriguingly, the model could barely account for the asymmetric traction-force loss at the end of the severed stress fiber (Figure 3f) and no traction-force loss occurred on the other side of the cell where the stress fiber was intact (Figure 3g). In addition, although we could define a given set of parameters for network elasticity that could account qualitatively for the various trends of traction-force relaxation, the traction-force changes were only in limited quantitative agreement with the equivalent experimental measurements (Figure 3b, d, f, g). Puzzled by the discrepancy between the predictions of the model and the measured loss of traction force on the other half of the cell after fiber photoablation, we decided to further interrogate the mechanical nature of the cortical meshwork.

### The cortical meshwork is contractile

In the experimental set up, shaving alone of a stress fiber, i.e. the longitudinal photoablation of the cortical meshwork, led to a significant release of contractile energy (Figure 4a and b). This release was comparable to that of fiber ablation. This result showed that the cortical meshwork was contractile and not passively elastic as initially hypothesized. It also meant that the cortical meshwork actively participated in the production of traction forces. Indeed, the release of mechanical energy after a single ablation and shaving of the stress fiber were additive (Figure 4a, b and Supplementary Movie S4), and consistent with the theoretical expectation these photoablations disrupted half of the contractile network (Figure 4a).

**Figure 4.**
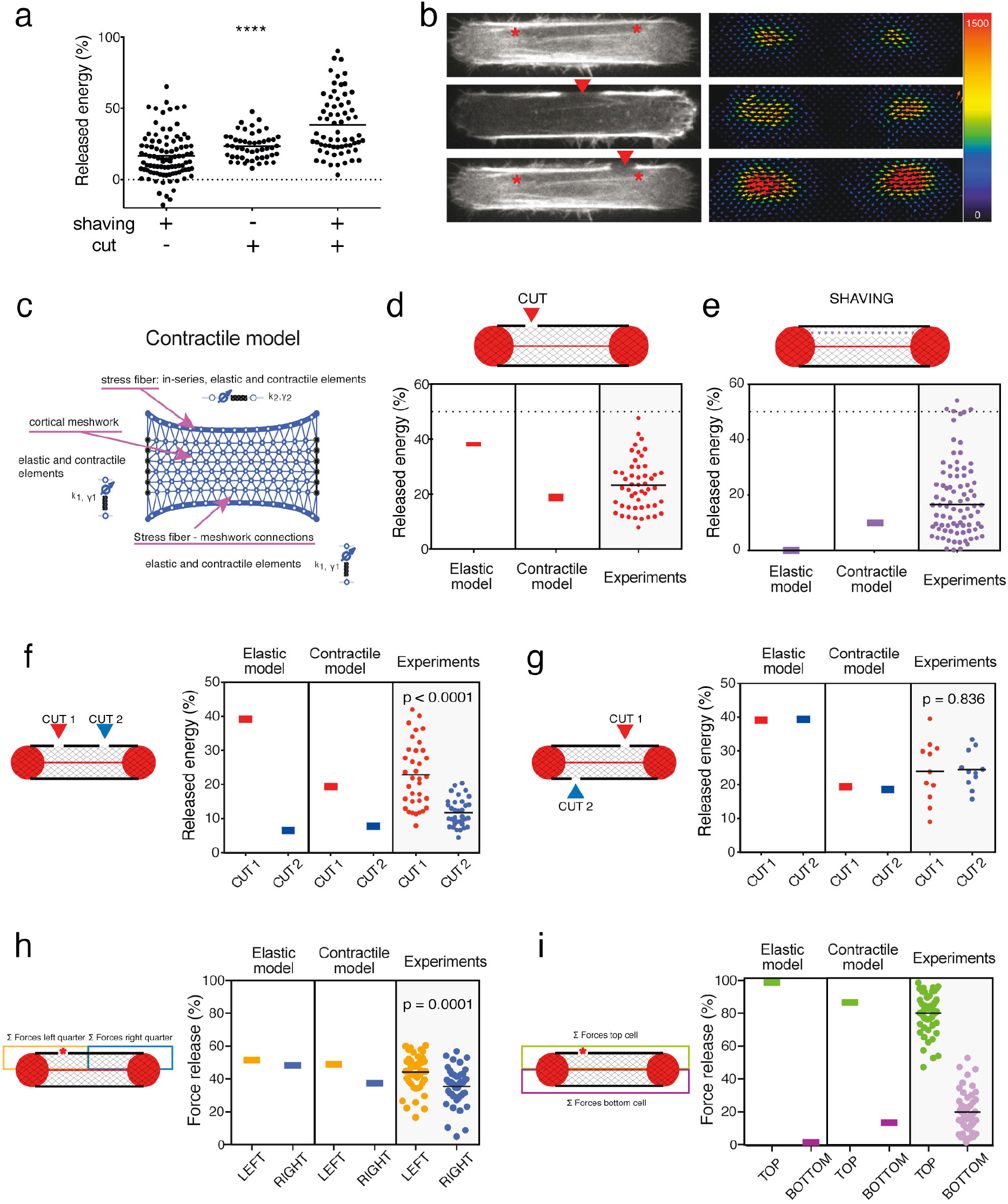
The cortical meshwork is contractile. a. Scatter plot of the released mechanical energy (percentage of the total mechanical energy before photoablation) after shaving the stress fiber, i.e. the photoablation of a narrow region medial and parallel to the length of the fiber above the non-adhesive substrate on the hydrogel, a single photoablation (cut) of the stress fiber, or a photoablation of the stress fiber after its shaving (shaving+cut). The p-value from a one way ANOVA test is indicated on the plot. b. From top to bottom: Images of RPE1-LifeAct-GFP cells on the left panel depicting a stress fiber subject to shaving, a single photoablation (cut), and shaving plus photoablation; with the corresponding traction-force maps of the cells shown in the right panel. The same cell was represented to illustrate the shaving and shaving plus photoablation (top and bottom panels, respectively). Image scale bar = 10 μm. Force scale bar in Pa. c. Diagram illustrating all components in the fully contractile model, including an isotropic contractile cortical mesh. d. Mechanical energy released after stress fiber photoablation (red arrow). Plot displaying the predictions of the elastic model, the contractile model and the experimental measurements (n=50 cells). e. Mechanical energy release after shaving (purple line). Plot displaying the predictions of the elastic model, the contractile model and the experimental measurements (n=95 cells). f. Sequential photoablations (cuts) on the same stress fiber at distinct locations (Cut 1=red arrow; Cut 2=blue arrow). Predictions of the elastic model, the contractile model and experimental data (n=35 cells). The p-value from a paired t-test is indicated on the plot. g. Sequential photoablations (cuts) on the two stress fibers in the cell (Cut 1=red arrow; Cut 2=blue arrow). Predictions of the elastic model, the contractile model and experimental data (n=11 cells). The p-value from a paired t-test is indicated on the plot. h. Spatial distribution of force loss along the stress fiber after off-center stress-fiber photoablation (red star). The loss of traction forces was considered in partitioned zones of the cell, where the orange zone included half the stress fiber and the off-centered photoablation site, and the blue zone included the other half of the stress fiber. Plot displaying the prediction of the elastic model, the contractile model, and the experimental data (n=47 cells). The p-value from a paired Wilcoxon test is indicated on the plot. i. Spatial distribution of force loss after off-center stress-fiber photoablation (red star). The loss of traction forces was considered in partitioned zones of the cell, where the green zone included the stress fiber with photoablation site, and the purple zone included the stress fiber without photoablation. Plot displaying the prediction of the elastic model, the contractile model, and the experimental data (n=47 cells).

Hence, we revised our initial *elastic* model by including contractility as a function of the cortical mesh (Figure 4c). The links in the cortical mesh were thus considered as contractile cables; i.e. tensed elastic springs in series with contractile elements (Figure 4c). In this second *contractile* model, the spring constants and contractilities of the cortical mesh and fiber-mesh links were the same, and different from those characterizing the stress fibers. The estimations of the magnitude and localization of mechanical-energy release based on this contractile model were in better agreement with the experimental data; including that the contractile energy released after stress fiber ablation was lower than in the elastic model, (Figure 4d), and the contractile energy released after stress fiber shaving was higher than in the elastic model (Figure 4e). As with the elastic model, the contractile model accounted correctly for the difference in contractile-energy release after the second photoablation whether it was applied to the same or opposite stress fiber. However, the amount of contractile-energy released in these scenarios was in better agreement with experimental data with the contractile model than with the elastic model (Figure 4f and g). More importantly, the contractile model captured key features of the spatial distribution of traction-force loss in response to a single photoablation in contrast to the elastic model. The contractile model recapitulated the asymmetry of traction-force loss at both ends of the severed fibers (Figure 4h) and the significant traction-force loss registered on the opposite side of the cell (Figure 4i). In addition, with the scenario of shaving the stress fiber prior to an off-center photoablation of that fiber (Supplementary Figure 3a), the contractile model, in comparison with the elastic model, better accounted for the left-right symmetrical loss of traction force (Supplementary Figure 3b; see also Figure 3f), and captured the loss of traction force at the opposite side of the cell (the side with the intact stress fiber; Supplementary Figure 3c).

The theoretical modelling combined with the experimental observations supported the hypothesis that the stress fibers and the cortical meshwork were mechanically similar in that they were both elastic and contractile. Therefore the stress fibers and the cortical meshwork may not be distinct networks with discrete interconnections, but part of a single integrated network, in which architecture and mechanics of actomyosin arrays vary in space.

### The cortical meshwork forms a continuum with stress fibers

To investigate further the similarities and differences in the structure and composition of the cortical meshwork and the stress fibers, immunostained actin filaments and myosin were visualized in fixed cells to a resolution of single actin filament by stochastic optical reconstruction microscopy (STORM). Instead of two distinct networks, the imaging revealed a continuous network, where filaments formed bundles which were progressively denser and more longitudinally-aligned with closer proximity to the peripheral stress fiber (Figure 5a). With high-resolution confocal microscopy, although the density of actin filaments was lower in the region between the two stress fibers and appeared darker compared to the high intensity of the two fibers, a meshwork of bundles and unresolved filaments could be visualized at higher signal saturation, together with numerous patches of myosin (Figure 5b). Indeed, myosins were densely packed along actin bundles (Figure 5b, red arrow heads), but were also present in regions in which no clear structure could be visualized (Figure 5b, red box). These observations confirmed that although their architectures were different, the stress fibers and the central mesh had a similar molecular composition.

**Figure 5.**
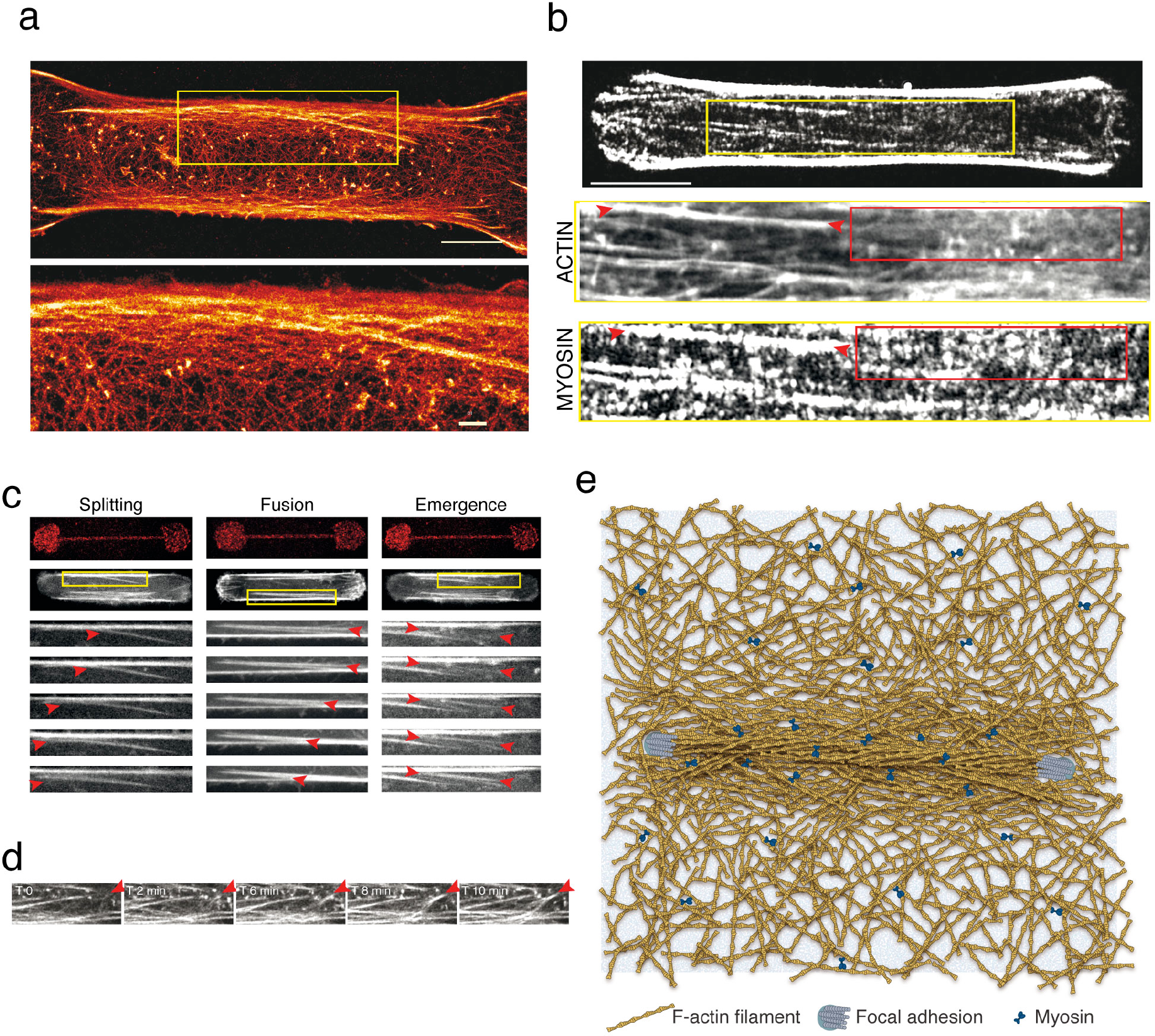
The stress fiber is fully embedded in the adjacent actin cortex and forms with it a continuous contractile network. a. STORM reconstructed image of the actin network (phalloidin-Alexa Fluor 647) in a RPE1 cell plated on a dumbbell-shaped micropattern on a glass substrate (scale bar=5 μm). The interconnectedness of the peripheral stress fiber with the surrounding actin cortex was highlighted in the yellow inset and associated zoom-in below (scale bar=1 μm). b. From top to bottom: Immunostainings of p-MLC (CY3) in a RPE1 cell spread on polyacrylamide dumbbell-shaped micropattern. Scale bar = 10 μm. Zoom-in images of the yellow inset are displayed below for actin (phalloidin-ATO-488) and p-MLC. For the p-MLC, signal was displayed at saturation in order to highlight small myosin patches inside the actin mesh. Red arrows indicate the area where actin structures are organized into bundles and red rectangles demark areas devoid of actin-identifiable structures. c. Live imaging of RPE1-LifeAct-GFP cells on dumbbell-shaped micropattern revealing various type of exchange between the peripheral stress fibers and bundles in the adjacent cortical meshwork. Images from top to bottom show sequential acquisitions with a time frame of 10 minutes. d. Live imaging of RPE1-LifeAct-GFP cells on dumbbell-shaped micropattern revealing the emergence (at the red arrow head) and connection of cortical bundles with peripheral stress fibers. e. Schematic representation of the stress fiber anchored at its two edges on the substrate via focal adhesions (blue disks) as a fully embedded structure within the surrounding contractile actin cortex (myosins are represented by blue bow-ties).

Live imaging of this network corroborated previous observations that cytoplasmic bundles permanently fused with or split from the peripheral fibers (Chen et al., 2019; Hirata et al., 2007; Hotulainen and Lappalainen, 2006; Luo et al., 2013; Muller et al., 2019) (Figure 5c, left and middle panels). Furthermore, cytoplasmic bundles were found capable to emerge from the cortical meshwork (Figure 5c, right, note the appearance of a bundle in between the arrow heads). Filaments in the cortical meshwork seemed to align and coalesce to form these bundles. The emerging bundle appeared capable to elongate by recruiting more cytoplasmic filaments and thereby connect to preexisting adjacent stress fibers (Figure 5d). This showed that stress fibers were not only connected to the meshwork but could also stem from the fusion with newly formed bundles in the meshwork; an assembly process that further accounts for their complete embedding in this meshwork. Interestingly, cortical bundles appeared more static above an adhesive area than above a non-adhesive area (Supplementary Figure 4a). This further suggested that the capacity to glide in the membrane helped the coalescence of thin bundles into larger stress fibers, whereas the presence of anchorage to the underlying extra-cellular matrix kept bundles separated. This absence of coalescence of small bundles into larger fibers above adhesive regions could account for lower energy release induced by bundle ablation in pill-shaped as compared to dumbell-shaped micropatterns (Supplementary Figure 4b,c) and for the difference in total cellular force production in those two geometries (Figure1c,d). Altogether, these results confirmed that peripheral stress fibers and the cortical meshwork form a dynamic and continuous contractile network.

## CONCLUSION

Our investigation of the force production by different actin networks revealed unexpected properties of the intracellular actomyosin machinery that appeared essential to integrate and transmit forces at the level of the cell. We demonstrated that the stress fibers are fully embedded in a contractile cortical actin network, and are not independent structures or structures with only discrete connections to other stress fibers. This meshwork of stress fibers and cortical filaments form a mechanical continuum. Our conclusion is in agreement with the previous ultrastructural observations of the cortical-actin–network connections to stress fibers and the more recent electron-microscopy demonstration that these connections depend on filamin A (Kumar et al., 2019; Marek et al., 1982; Svitkina, 2018). It also fits well with high-resolution imaging showing lateral interactions of myofibrils, which support a long range self-organization of contractile structures (Hu et al., 2017). Hence we found that contractile forces generated by the stress fiber are not only manifested at stress fiber anchorage points but are also propagated across the entire cell via the cortical meshwork. Also, we demonstrated that the contraction of the cortical meshwork actively contributes to traction force transmission to focal adhesions, thereby impacting the overall magnitude of contractile energy of the cell.

The mechanical interdependence of stress fibers and the cortical meshwork was supported by visualization of the network architecture in that filaments in the cortical meshwork tended to align with closer proximity to the stress fiber. This suggested that the randomly orientated filaments in the cortical network were converted into bundles of aligned filaments the nearer they were to the stress fiber, perhaps in a self-amplifying mechanism in which the bundle reinforces the tension in the stress fiber. We suspect this interconversion mechanism between the thin and non-attached filaments of the cortical network and bundled filaments to be essential for the rapid modulation of the production of traction forces on the microenvironment of the cell, in response to geometrical and mechanical cues. Such transitions in actomyosin network architecture were proposed to power cell-shape changes, either during mitosis or as cells progress through differentiation (Chalut and Paluch, 2016). In future work, the identification of the parameters triggering the dynamic exchange of filaments between these various contractile structures and understanding how the actin pool is distributed among them should help elucidate the link between actin network remodeling and force production at the cellular level.

## MATERIAL AND METHODS

### Preparation of micropatterned polyacrylamide gels

The preparation of patterned polyacrylamide hydrogels was performed according to the Mask method previously described in (Vignaud et al., 2014). A quartz photomask was first cleaned through oxygenplasma (AST product, 300 W) for 3.5 min at 200 W. Areas containing the patterns were then incubated with 0.1 mg/ml PLL-g-PEG (JenKem Technology ZL187P072) in 10mM HEPES pH 7.4, for 30 min. After de-wetting, the mask was exposed under deep-UV for 5 min. Next, patterns on the mask were incubated with a mix of 10 μg/ml fibronectin (#F1141, Sigma) and 20 μg/ml fibrinogen-Alexa-Fluor-647 conjugate (#F35200, Invitrogen) in 100mM sodium bicarbonate buffer pH=8.4 for 30 min. A mix of acrylamide (8%) and bis-acrylamide solution (0.264%) (Sigma) corresponding to the experimental Young modulus of 34.8 kPa was degassed for approximately 30 min, mixed with 0.2 μm PLL-PEG coated fluorescent beads (Fluorosphere #F8810, Life Technologies) and sonicated before addition of APS and TEMED. 25 μl of that solution was added on the micropatterned photomask, covered with a silanized coverslip (Silane, #M6514, Sigma) and allowed to polymerize for 25 min before being gently detached in the presence of sodium bicarbonate buffer. Micropatterns were stored overnight in sodium bicarbonate buffer at 4°C before plating cells.

### AFM measurements of the Young’s modulus of acrylamide gels

Gel stiffness was measured through nano-indentation using an atomic force microscope (Bruker Nanoscope) mounted with silica-bead-tipped cantilevers (r(bead) = 2.5 μm, nominal spring constant 0.06 N m^−1^, Novascan Technologies). The sensitivity of the photodiode to cantilever deflection was determined by measuring the slope of a force-distance curve when pressing the cantilever onto a glass coverslip. The force constant of the cantilever was determined using the thermal-noise method included in the Bruker Nanoscope software. For each acrylamide/bis-acrylamide ratio used in the traction-force microscopy measurements, 27 force curves in 3 by 3 grids were acquired (2 μm spacing between points) at three different locations on the gels. Before and during indentation experiments, gels were kept in PBS. Stiffness values from force curves were obtained using the NanoScope Analysis software, correcting for baseline tilt using the linear fitting option with the Hertz model and a Poisson ratio of 0.48 on the indentation curve.

### Preparation of micropatterned glass slides

To increase the resolution of actin images, RPE-1 cells were grown on glass micropatterns prepared as previously described in (Azioune et al., 2010). Glass coverslips were spin-coated for 30 sec at 3000 rpm with adhesion promoter Ti-Prime (MicroChemicals), then heated for 5 min at 120°C and spin-coated again for 30 sec at 1000 rpm with 1% polystyrene in toluene (Sigma). Coverslips were then oxidized by plasma (FEMTO, Diener Electronics) (19 sec, 30 W) and incubated for 30 min with 0.1 mg/ml PLL-g-PEG (PLL20K-G35-PEG2K, JenKem) in 10 mM HEPES pH 7.4. Dried coverslips where next exposed to deep-UV (UVO cleaner, Jelight) through a photomask (Toppan) for 5 min. Coverslips were incubated for 30 min with 10 μg/ml fibronectin (Sigma) and 20 μg/ml fibrinogen-Alexa-Fluor-647 conjugate (Invitrogen) in PBS (phosphate buffered saline) after UV exposure.

### Cell culture

Human telomerase-immortalized retinal-pigmented epithelial cells (RPE1; Clontech) either expressing LifeAct-GFP or parental (Vignaud et al., 2012) were grown in a humidified incubator at 37°C and 5% CO_2_ in DMEM/F12 medium supplemented with 10% fetal bovine serum and 1% penicillin/streptomycin (GIBCO/Life technologies). Cells were plated at approximately 15000 cells/ml on patterned polyacrylamide gels and left to spread for 3 to 4 hours before imaging.

### Immunostaining and labeling

Cells were pre-permeabilized in 0.5% Triton X-100 in cytoskeleton buffer for 17 sec for p-MLC and alpha-actinin staining and then rapidly fixed in 4% paraformaldehyde in cytoskeleton buffer 10% sucrose pH 6.1 for 15 min at room temperature. Cells were then washed twice with cytoskeleton buffer and incubated in quenching agent 0.1 M ammonium chloride for 10 min.

For all conditions, after fixation, the cells were washed then blocked with 1.5% bovine serum albumin (BSA) for 45 minutes. The cells were incubated with appropriate dilutions of primary antibodies in PBS containing 1.5% BSA and 0.1% Tween overnight at 4°C in a humid chamber. For the primary antibodies, anti-phospho-myosin light chain 2 (#3671, CST), anti α-actinin (#05-384, Millipore), and anti-paxilin (#610051, BD Biosciences) were used. After several washing steps, the coverslips were then incubated with secondary antibodies (Alexa-Fluor antibodies, Invitrogen) diluted in PBS with 1.5% BSA and 0.1% Tween for 1 h at room temperature in a humid chamber. After washing, Phalloidin-FITC (#P5282, Sigma) was incubated for 20 min. After washing, coverslips were then mounted onto slides using Prolong Gold antifade reagent with DAPI (#P36935, Invitrogen). Fluorescent Tetraspeck microspheres of 0.5 μm diameter (#T7281, Life Technologies) were in some cases incubated with the coverslip to provide an internal fluorescence intensity reference. Whenever needed, SirActin (SC001, Spirochrome) was used at a concentration of 500 nM for 3 h to stain actin in living cells.

### STORM imaging

After fixation and immunolabeling, cells were incubated with phalloidin-Alexa-Fluor-647 (0.5 μM, Thermo Fisher) overnight at 4°C. After two quick rinses in phosphate buffer, RPE1 cells were mounted in a closed chamber in STORM buffer (Smart kit, Abbelight) and imaged by STORM as described previously (Ganguly et al., 2015; Jimenez et al., 2019) using an N-STORM microscope (Nikon Instruments) equipped with an Ixon DU-897 camera (Andor). Phalloidin (0.25μM) was added in the STORM medium to mitigate progressive unbinding from actin filaments during imaging (Jimenez et al., 2019). After locating a cell using low-intensity illumination, epifluorescence images were acquired in both the green and far-red channels. For STORM imaging of actin, the sample was continuously illuminated at 647nm (full power) and a series of 60000 to 100000 images (256×256 pixels, 15 ms exposure time). The N-STORM software (Nikon Instruments) was used for the localization of single fluorophore activations. After filtering, localizations with more than 800 photons, the list of localizations was exported as a text file and the ThunderSTORM plugin (Ovesný et al., 2014) of Fiji was used to generate reconstructions.

### Image acquisition and photoablation

Images of the different immunostainings and high-resolution time-lapse of actin dynamics on PILL and DUMBBELL micropatterns were acquired on a confocal microscope Zeiss LSM800 using a 63X maginification objective. Staining of p-MLC, α-actinin and paxilin were imaged using an AiryScan detector. GaAsP detectors were used for DAPI and micropattern stainings.

Traction-force mapping, together with regular RPE1-LA-GFP, actin-GFP (CellLight™ Actin-GFP, BacMam 2.0 from ThermoFischer Scientific) or SirActin (SC001, Spirochrome) imaging, were performed on a confocal spinning-disk system (EclipseTi-E Nikon inverted microscope equipped with a CSUX1-A1 Yokogawa confocal head, an Evolve EMCCD camera from Roper Scientific, Princeton Instruments).

Photoablation was performed on a spinning-disk system from Nikon using the iLas2 device (Gataca Systems) equipped with a passively Q-switched laser (STV-E, ReamPhotonics, France) at 355 nm producing 500 picoseconds pulses. Laser displacement, exposure time and repetition rate were controlled via ILas software interfaced with MetaMorph (Universal Imaging Corporation). Laser photoablation and subsequent imaging was performed with a 100X CFI S Fluor oil objective (MRH02900, Nikon) or a 60X CFI S PLAN FLUOR ELWD objective. The stress-fiber punctual photoablation was performed on fly during live acquisition. For the stress-fiber shaving, photoablation was performed on a narrow region medial and parallel to the length of the fiber above the non-adhesive substrate on the hydrogel. For all photoablations, 13 repetitions of 25 ms pulses were used with 100% of the 355nm laser power, corresponding to a pulse of approximately 450 ms.

### Measurement of cell traction forces with ImageJ

Data were analysed with a set of macros in Fiji using the method previously described in (Martiel et al., 2015). Displacement fields were obtained from fluorescent bead images before and after removal of cells by trypsin treatment. Bead images were first paired and aligned to correct for experimental drift. Displacement field was calculated by particle imaging velocimetry (PIV) on the basis of normalized cross-correlation following an iterative scheme. Final vector-grid size ranged from 1.55 μm × 1.55 μm to 1.60 μm × 1.60 μm depending on magnification. Erroneous vectors where discarded owing to their low correlation value and replaced by the median value of the neighbouring vectors. Traction-force field was subsequently reconstructed by Fourier-transform traction cytometry, with a regularization parameter set to 8×10^−11^. Force vectors located outside of the micropattern area were discarded.

### Force quadrant analysis

Cell-traction force was computed above. The traction-force field was divided into 4 zones using the two planes of symmetry of the dumbbell shape of the micropattern. In each zone, forces were summed-up vectorially. The resulting vector was then located at the center of the zone for display.

### Statistical analysis

Statistical analysis and chart design was performed using Graphpad Prism 6 (www.graphpad.com) and R version 3.4.0 together with RStudio version 1.0.143.

## ACKNOWLEDGEMENTS

We thank the live microscopy facility MuLife of IRIG/DBSCI, funded by CEA Nanobio and labex Gral for equipment access and use.

