## Supplemental Information for "Stress fibers are embedded in a contractile cortical network"

### SUPPLEMENTARY INFORMATION

- Supplementary methods
- Supplementary references
- Supplementary figures
- Supplementary movies

### Supplementary method

MODEL DESCRIPTION

The actin cytoskeletal meshwork is modeled as a two-dimensional deformable elastic material (Copos et al., 2017; Guthardt Torres et al., 2012; Kohler and Bausch, 2012; Zhu and Mogilner, 2012; Besser et al., 2011; Bischofs et al., 2008). This is a simplification of a three-dimensional model since on average the height of an adherent cell is much smaller than its in-plane dimensions, and so we neglect any out-of-plane deformations. The rheology of the cytoskeleton meshwork has been described as a viscoelastic gel with time-evolving material properties due to the turnover of actin and action of molecular motors (Moeendarbary et al., 2013; Mofrad, 2009; Gardel et al., 2008). There are two possible theoretical descriptions used to capture this rheology: (a) elastic elements embedded in a viscous gel (Barnhart et al., 2015; Larripa and Mogilner, 2006), or (b) viscous elements embedded in an elastic gel (Strychalski et al., 2015; Zhu and Mogilner, 2012; Joanny and Prost, 2009). The loss of traction force due to an ablation suggests that on timescales of tens of seconds the material response is well described by an elastic gel with viscous elements (Figure 1f). Further, it has been reported that the viscous timescale of the actin gel due to the growth and reorganization of actin filaments (order of minutes) is much longer than our observation time (Moeendarbary et al., 2013; Gardel et al., 2008).

For simplicity, the elastic gel is coarse-grained into a network of nodes interconnected by elastic springs and viscous elements are excluded as their dynamics would only affect the transient behavior (Zhu and Mogilner, 2012; Zimmermann et al., 2010). An interconnected network of nodes and elastic links is a general theoretical approach used in previously published models to describe the mechanics of the cell interior of motile and adherent cells (Copos et al., 2017, Guthardt Torres et al., 2012; Zhu and Mogilner; 2012; Besser et al., 2011, Bischofs et al., 2008). While continuum approaches have been developed and successfully used as well (Barnhart et al., 2015; Oakes et al., 2014; Joanny and Prost, 2009), there is a tradition of modeling the cytoskeletal meshwork using discrete elements: nodes and springs (Boal, 2012).

Specifically, in the model, the cell interior is a rectangular network of nodes connected by Hookean elastic springs with endpoints which are adhering to the substrate following experimental evidence (Figure 3a). The nodes represent material points of the cortical meshwork while the elastic springs interconnecting the nodes model the mechanical response of the actin filament arrays. In the experiments with a dumbbell adhesive micropattern, large contractile actomyosin bundles are reported along the cell periphery stretching between the adhesive endpoints (Figure 1b). We include these bundles as additional nodes placed along a line at the top and bottom interface of the elastic meshwork. The nodes representing the actomyosin bundles are interconnected by Hookean elastic springs with an additional tension to mimic the effect of myosin-generated contractility. The nodes are also connected to the meshwork via additional linear elastic links. The endpoints of the contractile fibers are anchored to the substrate as was done with the endpoints of the elastic meshwork.

At the beginning of the simulation, the initial network of nodes and springs is the result of a Delaunay triangulation algorithm applied to a rectangular domain. The nodes are chosen to be the vertices of the triangulation and the elastic springs lie on the edges of the triangulation. All springs are created in an undeformed state. The linear springs in the meshwork are characterized by a stiffness $k_1$, while the additional links representing the actomyosin bundles have spring stiffness $k_2$ and tension $\gamma_2$. The connectors between the contractile fibers and the elastic meshwork have the same spring stiffness as the meshwork $k_1$.

Given this initial shape, elastic forces are computed at every node in the network. As was done previously (Copos et al., 2017; Barnhart et al., 2015), we assume that adhesion complexes generate viscous resistance to the deformation of the meshwork. The respective resistive force is given by $\xi \vec{u}$ where $\xi$ is the effective adhesion drag coefficient and $\vec{u}$ is the velocity of the network in the lab coordinate system. The adhesion resistive force is balanced by the active elastic stresses: $\xi \vec{u} = \vec{F}$ at every material point. At each time step, the velocity of a node is determined from the force balance. At the endpoints of the network, the velocity is enforced to be zero since the cell is adhering to the substrate at these locations. Once the position of the nodes is updated in time, elastic forces are recomputed at these new locations and the algorithm proceeds as indicated above. Because of the relatively small deformations we never observe instabilities or crossovers in the triangulation. When the network achieves mechanical equilibrium, the forces exerted by the meshwork on the substrate (i.e., at the endpoints) are recorded as traction forces and qualitatively compared with experiments.

The scope of this theoretical framework is to probe the respective contribution of actomyosin bundled stress fibers, the internal cortical meshwork, and their mechanical coupling on exerted traction forces. The investigation is achieved by qualitative comparison of model predictions of the spatiotemporal distribution of traction forces and experimental measurements from adherent cells. For computations, we normalize the parameters so that $k_1 = 1$ for the stiffness of the elastic network and the stiffness of the connections between the elastic meshwork and the contractile fibers. For the contractile fibers we choose a larger spring stiffness of $k_2 = 20$ and tension $\gamma_2 = 2$ relative to the normalized stiffness of the elastic mesh. The value for the spring stiffness $k_2$ is chosen to qualitatively reproduce the force loss for two consecutive cuts along the same fiber; specifically, the difference in the force released between the two cuts would be less significant by increasing $k_2$. In the modified model with a contractile cortical meshwork, we found that decreasing $k_2$ results in too much force loss in the region the cell opposite of the injured stress fiber. The value for the tension in the stress fiber, $\gamma_2$, is arbitrarily chosen, but its value relative to $\gamma_1$ is important. First, indifferent of our numeric choices for the stress fiber parameters, $k_2$ and $\gamma_2$, in the absence of a tension in the meshwork (i.e., $\gamma_1 = 0$) ablation of the mesh would never produce a force loss (Figure 3g). Second, introducing too large of a tension in the meshwork would imply that cuts along the fiber release very little force. To introduce cortical tension, we set $\gamma_1 = 0.5 = 0.25\gamma_2$ to simultaneously reproduce the force loss due to a single cut along a stress fiber and force loss due to an ablation of the cortical network. In particular, there are two competing effects that limit the choice of $\gamma_1$ — increasing $\gamma_1$ increases the force released with ablation since the cortical mesh participates more in generating contraction, but increasing $\gamma_1$ decreases the force released with a single cut on the fiber. Each parameter, $k_2, \gamma_2$, is varied individually tenfold and chosen to match as close as possible the qualitative behavior seen in the experiments. The drag coefficient is 0.025 arbitrary force per velocity units (2 orders of magnitude smaller than the elastic forces). Because traction forces are computed after the meshwork relaxes to the equilibrium, the value of the drag coefficient does not affect our results. The timestep is $\Delta t = 0.0001$ arbitrary time units and is chosen to meet numerical stability constraints.

### Supplementary figures

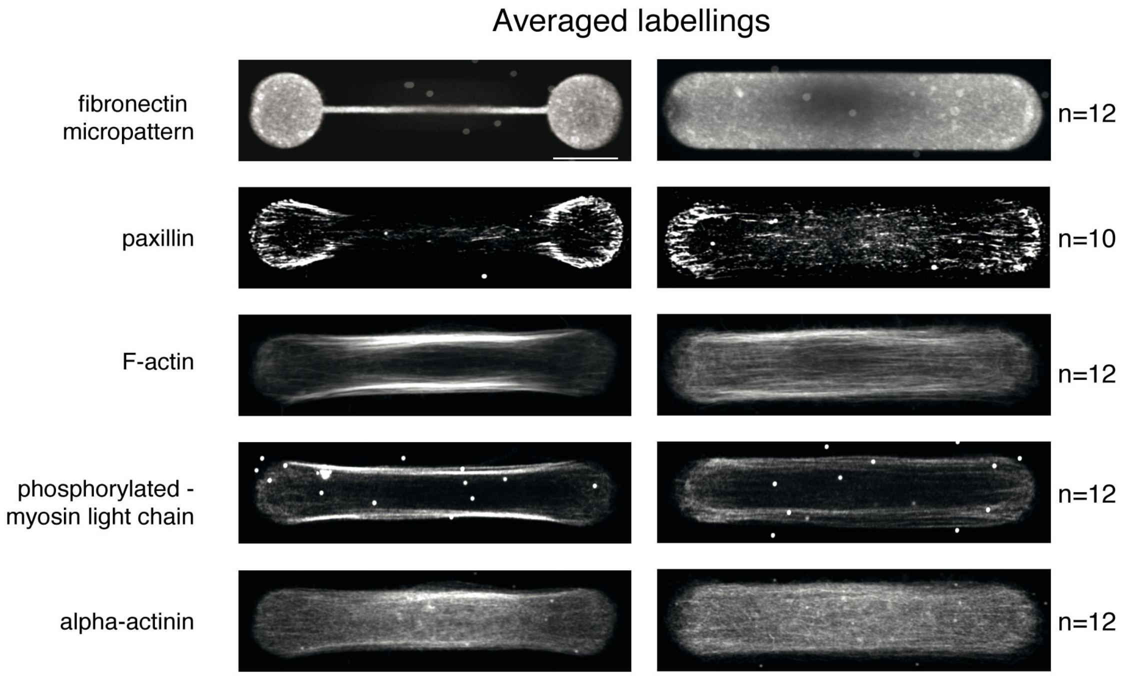


**Supplementary Figure 1 - Actin networks composition.**

Averaged distribution of molecular components involved in cell contractility for cells displaying either two main peripheral stress fibers or a continuous actin mesh.
Immunostainings of RPE1 cells spread respectively on dumbbell (left panel) and on pill micropatterns (right panel). For each shape, averaged Z projections of cells are displayed. From top to bottom: micropatterns labeling (fibrinogen-Cy5); actin (phalloidin-ATO-488); paxillin (alexa-488); phospho-MLC (CY3); alpha-actinin (CY3). Image scale bar = 10 µm.

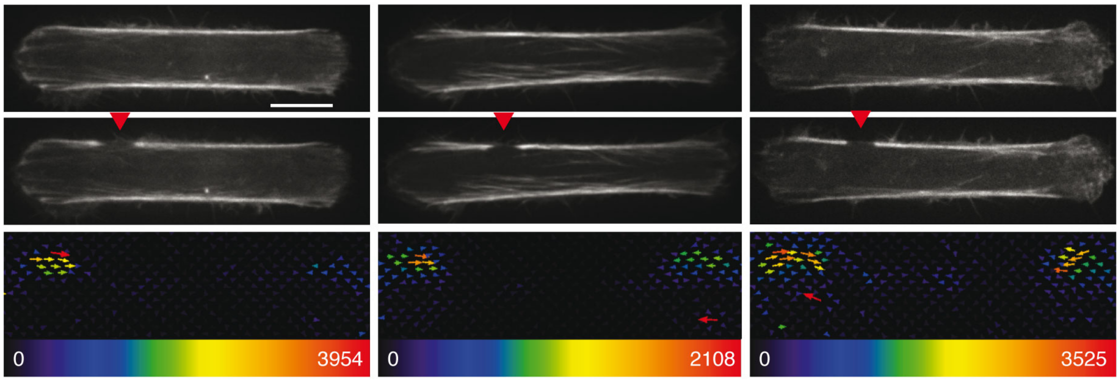


**Supplementary Figure 2 - Asymmetric force relaxation after an off-centered stress fiber cut.**

Images of RPE1-LA-GFP cells after off-centered photoablation of the stress fiber (red arrow) and the associated traction maps. Image scale bar = 10 µm. Force scale bar in Pa.

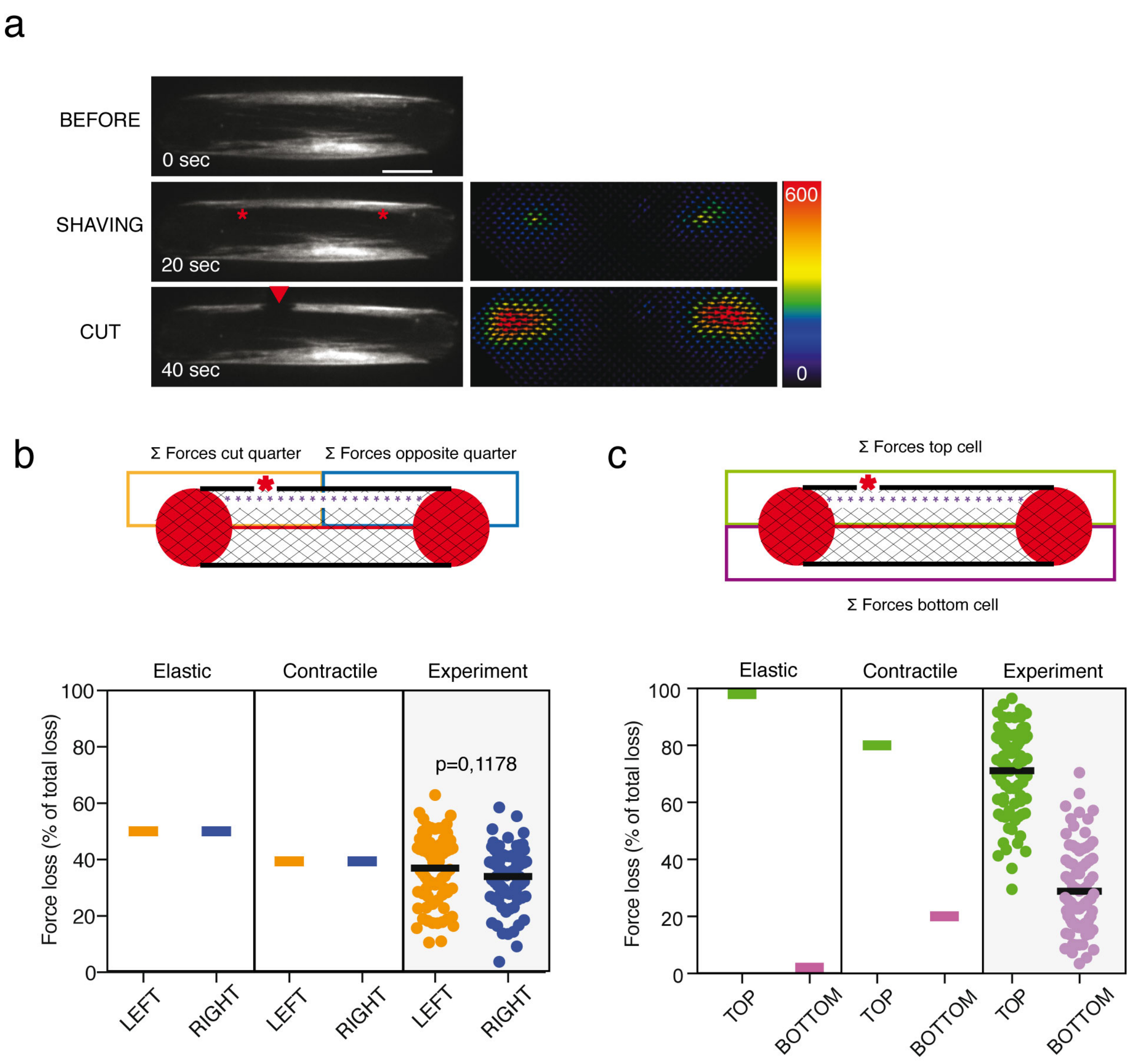


**Supplementary Figure 3 - Elastic and contractile model predictions of the force loss distribution after stress fiber shaving and photoablation.**

a. From top to bottom: Images of a representative RPE1-LA-GFP cell depicting the precut cell; a shaving; a consecutive cut together with the associated forces relaxed upon photoablation on the right panel. Image scale bar = 10 µm. Force scale bar in Pascal.
b. Spatial distribution of force loss along the stress fiber after stress-fiber shaving (purple dashed line) and off-center photoablation (red star). The loss of traction forces was considered in partitioned zones of the cell, where the orange zone included half the stress fiber and the off-centered photoablation site, and the blue zone included the other half of the stress fiber. Plot displaying the predictions of the elastic model, the contractile model and the experimental measurements (n=80 cells). The p-value from a paired t-test is indicated on the plot.

c. Spatial distribution of force loss along the stress fiber after stress-fiber shaving (purple dashed line) and off-center photoablation (red star). The loss of traction forces was considered in partitioned zones of the cell, where the green zone included the stress fiber with photoablation site, and the purple zone included the stress fiber without photoablation. Plot displaying the predictions of the elastic model, the contractile model and the experimental measurements (n=80 cells).

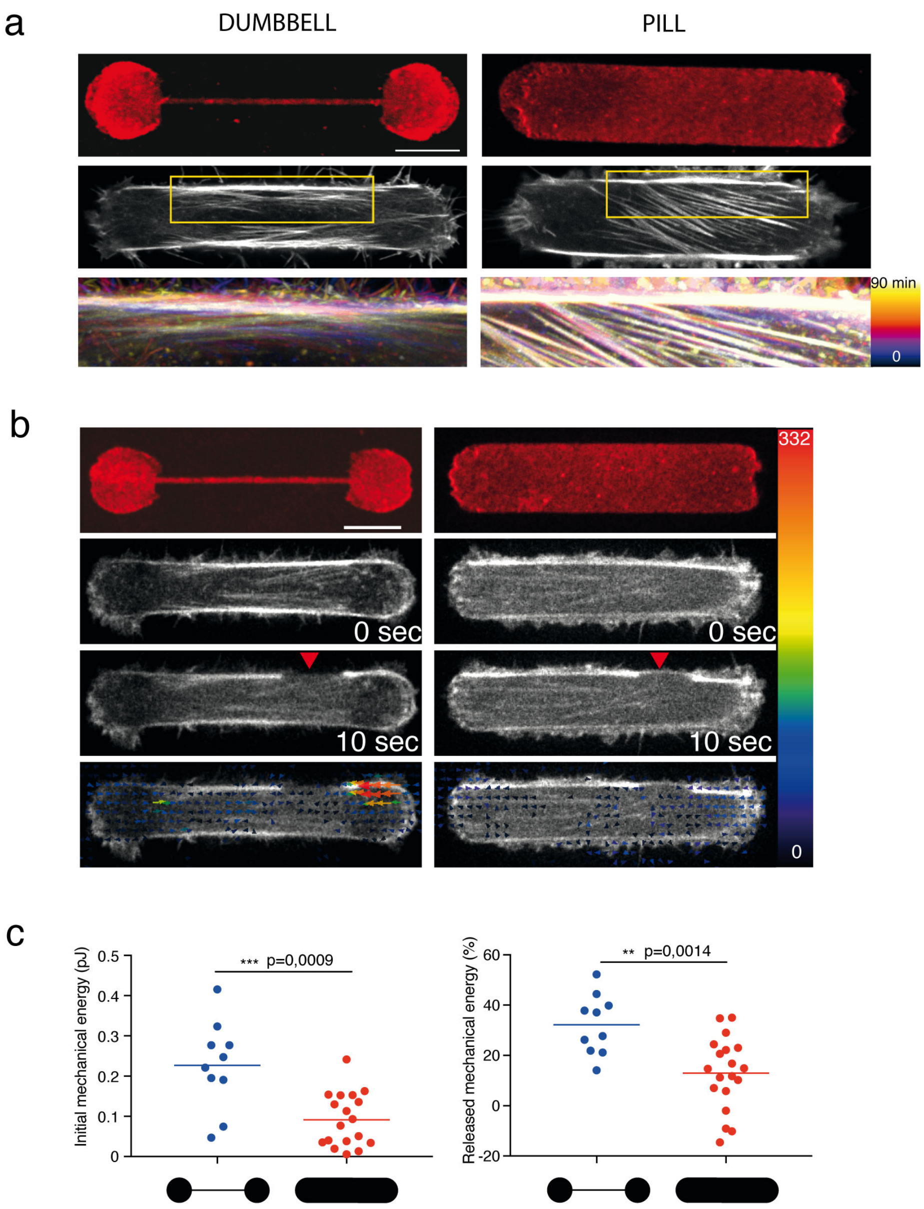


**Supplementary Figure 4 – Dynamics and mechanics of cortical bundle fusion with lateral stress fibers depending on anchorage with the underlying extra-cellular matrix.**

a. Color-coded overlay of sequential images acquired every 10 minutes on dumbbell-shaped (left) and pill-shaped micropatterns (right). Color overlay showed static structures in white and moving structures in colors ranging from blue to yellow depending on the time frame in which they were acquired.
b. From top to bottom: Micropattern labeling (fibrinogen-CY5); Actin before stress-fiber photoablation (0 sec); Actin (LA-GFP) after stress-fiber photoablation (10 sec); Overlay

of the traction-force map and actin-staining image after photoablation. Image scale bar = 10 µm.

c. Left panel: Scatter plot of the initial mechanical energy of the cells before the photoablation. Right panel: Scatter plot of the released mechanical energy following photoablation of the peripheral stress fiber (% of the initial mechanical energy). N=1, n= 10 for dumbbell and n=18 for pill. The p-values from a Mann-Whitney t-test are indicated on the plots.

#### Supplementary movies

Supplementary Movie S1

Illustrations of stress fiber ablation (showed with white arrow heads) in nine cells plated on dumbbell-shaped micropatterns on poly-acrylamide gels. Scale bar is 10 μm.

Supplementary Movie S2

Illustration of stress fiber ablation and the measurement of the corresponding traction force field relaxation in a cells plated on dumbbell-shaped micropatterns on poly-acrylamide gel. Scale bar is 10 μm.

Supplementary Movie S3

Illustration of stress fiber shaving (in between the two tilted arrow heads) and the measurement of the corresponding traction force field relaxation in a cells plated on dumbbell-shaped micropatterns on poly-acrylamide gel. Scale bar is 10 μm.

Supplementary Movie S4

Illustrations of stress fiber shaving (in between the two tilted arrow heads) followed by stress fiber ablation (single vertical arrow head) and the measurement of the corresponding traction force field relaxation in a cells plated on dumbbell-shaped micropatterns on poly-acrylamide gel. Scale bar is 10 μm.
